# Bioelectrical pattern discrimination of *Miconia* plants by spectral analysis and machine learning

**DOI:** 10.1101/2020.11.10.377036

**Authors:** Valéria M M Gimenez, Patrícia M Pauletti, Ana Carolina Sousa Silva, Ernane José Xavier Costa

## Abstract

We have conducted an *in loco* investigation into the species *Miconia albicans* (SW.) Triana and *Miconia chamissois* Naudin (Melastomataceae), distributed in different phytophysiognomies of three Cerrado fragments in the State of São Paulo, Brazil, to characterize their oscillatory bioelectrical signals and to find out whether these signals have distinct spectral density. The experiments provided a sample bank of bioelectrical amplitudes, which were analyzed in the time and frequency domain. On the basis of the power spectral density (PSD) and machine learning techniques, analyses in the frequency domain suggested that each species has a characteristic biological pattern. Comparison between the oscillatory behavior of the species clearly showed that they have bioelectrical features, that collecting data is feasible, that *Miconia* display a bioelectrical pattern, and that environmental factors influence this pattern. From the point of view of experimental Botany, new questions and concepts must be formulated to advance understanding of the interactions between the communicative nature of plants and the environment. The results of this on-site technique represent a new methodology to acquire non-invasive information that might be associated with physiological, chemical, and ecological aspects of plants.

**Highlight:** *In loco* characterization of the bioelectrical signals of two *Miconia* species in the time and frequency domain suggests that the species have distinct biological patterns.

## Introduction

The bioelectrical signals of plants can be classified into action potential (AP), variation potential (VP), systemic potential (SP), local electrical potential (LEP) (Fromm and Lautner, 2007), and oscillatory signal (OS) (Cabral et al., 2011). OS can be understood as a sum of all the other signals. OS can be acquired by invasive (electrode-intracellular needle) or non-invasive (disc-extracellular electrode) techniques. To characterize OS, visually distinct low-amplitude VP variations must be recorded. These variations show how OS behaves as a function of time and frequency. Oscillations occur under different conditions, thereby generating analyzable spectra (Cabral et al., 2011).

Thus, apparently immobile plants are in fact extremely dynamic beings. They are sensitive to environmental changes that may limit their chances of development and survival (Vodeneev, Akinchits, and Sukhov, 2015). Plants are subject to stimuli that trigger the transmission of information through signaling systems. These systems consist of networks that are integrated by connections between their elements, which are affected by hydraulic, chemical, or electrical action (Trewavas, 2007). After an excitation threshold is reached, stimulation produces an AP that is propagated to other plant cells and which may be associated with inhibition or expression of genes related to tolerance to various types of stress. APs are identified as electrical messages. In animals, APs take place in axons, where the ionic mechanism depends on Na^+^ ion flow (inward depolarization) and K^+^ ion output (repolarization) (Fromm and Lautner, 2007). In plant, AP generation and transmission imply changes in ion flow, such as Cl^−^ and K^+^ efflux and Ca^2+^ inflow. This causes an ionic current and evidences that this mechanism is associated with water efflux and turgor loss (Volkov and Markin, 2015).

Plant electrical signals, which are mediated by cytosolic Ca^2+^ kinetics at the cell membrane level, usually generate a response within less than one second (Trewavas and Baluška, 2011). Therefore, electrical signals are crucial to plants. The speed at which information is transmitted distinguishes such signals from other forms of signaling and elicits ultrafast responses coordinated by appropriate cells (Davies, 2006; Fromm and Lautner, 2007; de Toledo et al., 2019). Over long distances, such signals are affected by chemical signals until they reach the vascular system or change gene expression, which can culminate in increased circulation, synthesis, and emission of volatile compounds like jasmonic acid and ethylenes into the atmosphere (Baluška and Mancuso, 2009; Volkov and Marquin, 2015). Volatile signals constitute local defenses and can be detected by weakly connected areas of the plant and by surrounding plants for many days (Baluška and Mancuso, 2009; Ali et al., 2013; Volkov and Markin, 2015). Mechanisms for converting electrical signals into functional responses have also been investigated (Vodeneev, Katicheva, and Sukhov, 2016). Some results have suggested a communication model based on AP transmission along highly reticulated vascular conduits (Dinant and Lemoine, 2010; Calvo, 2016; Calvo, Sahi, and Trewavas, 2017).

Studies have shown that the electrical potential of plants expands the possibilities of recognizing and classifying information fast in response to specific environmental stimuli (Volkov, 2012; Huber and Bauerle, 2016; Chen et al., 2016; Calvo and Friston, 2017). In this way, physiological and morphological aspects can be grouped into responses to these circumstances, indicating that plants prioritize signals among the several sources of stimulation (Calvo and Friston, 2017). Plants can still adapt to environmental fluctuations by generating complex systems that can withstand these variations with greater stability (Souza, Pincus, and Monteiro 2005; Volkov and Markin, 2015; Volkov et al., 2017; Pereira et al., 2018). Therefore, electrical signals are physical entities that can carry information about the nature or behavior of a given phenomenon and which vary over time and space (de Toledo et al., 2019), so that plants can be considered a network of transducers that receive and emit different types of signals. The electrochemical-biochemical signal plays an important role in the communication between internal components associated with biotic or abiotic environmental conditions, allowing the physiological activities of plants to be characterized (Hasegawa et al., 2000).

Additionally, computer technology and mathematical models have evolved, allowing a plant’s electrical signal to be collected, interpreted, encoded, and transformed into a new “language” that can reflect plant physiology and the interaction of the plant with the environment at the individual level. This helps to understand the principles and relationships that govern the plant’s organizational behavior and suggests bioelectrical models (Souza et al., 2017).

In the context of the omic sciences, electroma means the totality of all ionic currents of any living entity from the cellular to the organismal level that is vital for the maintenance and definition of life, whereas cellular “death” is the moment when the ability to perceive the electrical dimension is irreversibly (excluding regeneration) lost (De Loof, 2016).

Bioelectrical signals obtained from plants with amplifier equipment generate large datasets that can be processed by digital signal processing and machine learning techniques. Power spectral density (PSD) estimation is a relevant digital signal processing (DSP) technique that can be used to analyze periodic and random signals (Proakis and Manolakis, 1996). Numerous problems in signals acquired from biological systems can be tackled by applying spectrum analysis as a preliminary measurement before further processing is performed (Fong et al., 2016). Sophisticated spectrum measurements can be used to obtain information from bioelectrical signals; for example, EEG (Manshouri, Maleki, and Kayikcioglu, 2018; Costa and Cabral, 2000) and ECG (Baldin et al., 2020).

Spectrum analysis by PSD encompasses various measurements. For this reason, this topic has received close attention and has been employed in several research areas. Moreover, applying a machine learning algorithm like Random Forests (RF) is useful when such extensive arrays of data are analyzed (Verikas, Gelzinis, and Bacauskiene, 2011). RF is a bootstrapping classification tool that generates decision trees by using different sets of randomly selected input variables, to make a prediction. The RF algorithm was introduced by Breiman Leo in 2001 (Breiman Leo, 2001), and it can be employed for classification and regression. It can be successfully applied in statistical analysis because it is not complex, has few parameters to tune, and can tackle high-dimensional feature spaces and complex data structures (Oparin et al., 2008; Hernandez et al., 2008).

The accumulated knowledge about different electrical signals in plants and their relationship with short- and long-distance transmission of physiological responses has been established in a controlled environment, mainly for monitoring physiological aspects related to plant development and for understanding agronomic factors related to production. However, this resource has not been used to understand plant behavior *in loco*. In this scenario, this study aimed to collect the oscillatory bioelectrical signals of two Brazilian Cerrado species of *Miconia* (Melastomataceae) *in loco* and to verify if the patterns of the signals differ in terms of spectral density.

## Materials and methods

### Study Areas

The bioelectrical signals were collected in three fragments of Cerrado in different locations of the state of São Paulo. Two of these fragments are part of conservation units: Estação Ecológica de Jataí (EEJ), in Luiz Antônio (21°33’ S and 47°51’ W), and Parque Estadual Furnas do Bom Jesus (P.E.F.B.J.), in Pedregulho (20°08’ S and 47°16’ W). The third fragment is located at the University of São Paulo in Pirassununga (21°36’ S and 47°15’ W). These fragments have distinct characteristics.

According to the Köpen-Geiger climatic classification, experimental area 1 (EA1, Pirassununga) presents Cwa climate, which represents humid temperate climate with dry winter and hot summer; experimental area 2 (EA2, Luiz Antônio) is classified as Aw, corresponding to tropical climate with dry winter season; and experimental area 3 (EA3, Pedregulho) corresponds to Cwb, where the climate is temperate humid with dry winter and temperate summer.

In each of the three study areas, plots of 40 m × 50 m were established in two Cerrado phytophysiognomies: Cerrado *stricto sensu* (dry environment), where seven Ma (*Miconia albicans*) specimens were sampled, and gallery forests or areas close to a watercourse (wet environment), where seven Ma specimens and seven Mc (*Miconia chamissois*) specimens were sampled. Thus, 21 individuals were sampled in each study area, to give a total of 63 individuals.

In EA1, the plot located in Cerrado *stricto sensu* was close to vegetation in an advanced stage of conservation. The Ma specimens consisted of adult plants measuring up to 3 m that were sparsely distributed among dense vegetation of large size (4 to 8 m) and which relied on abundant shade and dry soil; winds and grasses were absent. In the same study area, Mc was found in a plot located on the edge of a lagoon (Pindaíba), where the soil was soaked, the density of the vegetation provided different shade, humidity, and temperature than the plot located in Cerrado *stricto sensu*.

In EA2, the plot located in Cerrado *stricto sensu* was delimited and was undergoing regeneration. The vegetation measured between 3 and 6 m, on average, and the canopy was discontinuous due to the distance between the plants. Light incidence was high, and the soil was dry and sandy. The Ma population density in this environment was high; the population was dispersed in the area and was becoming aggregated at the edges of the fragment. In this same study area, the other plot was located on the banks of the Bandeira stream, where the larger vegetation (4 to 10 m) was dense and shaded. The soil was dark and moist and aggregated extensive Mc populations, as well as some Ma specimens growing in areas where the soil was drier.

EA3 was characterized by the dominant presence of preserved Cerrado *stricto sensu*, with vegetation measuring between 3 and 6 m, on average. The canopy was discontinuous, and light and wind incidence among the plants was high. The soil was rocky. The presence of grasses was intense among the vegetation where Ma occurred with sparse distribution. Mc was found as populations aggregated inside large shaded depressions (furnas) where the moist soil became soaked in some stretches. In the “furnas”, the plants were protected from wind. At the high and sunny edges, the soil was dry, and Ma was sparse.

### Botanical Materials

Among Angiosperms, Melastomataceae Jussieu is the seventh most diverse and the second most common botanical family. It has pantropical distribution and high neotropic concentration, and it comprises about 166 genera and 4,570 species including herbs, shrubs, and trees. Thirty genera and 248 species occur in the state of São Paulo (Wanderley et al. 2009; Oliveira and Marquis, 2002). However, the conservation status of this family is worrisome because habitats that aggregate a high number of endemic species, with restricted and punctual distribution, are being destroyed (Morandi et al., 2020).

*Miconia* is the largest genus in terms of the number of species—approximately 1,050 *Miconia* species exist. In Brazil, there are 289 *Miconia* species, 122 of which are endemic (Zappi et al., 2015). Two species were selected for this study. One of them is *Miconia albicans* (SW.) Triana, which is distributed in South America and southern Mexico, is considered a pioneer species, and occurs in the fields of Cerrado, the Amazon Forest, Caatinga, and the Atlantic Forest (Neri et al., 2005; Allenspach and Dias, 2012). It is a shrub or small tree reaching up to 3 m in height, and it is popularly known as “pixirica” or “folha branca”. In the state of São Paulo, this species can be found in forest edges of typical phytophysiognomies or in Cerrado regenerated areas (Durigan et al. 2004). The other species that was selected for this study is *Miconia chamissois* Naudin, which occurs from Mexico to southern Brazil and Argentina. *M. chamissois* is popularly known as “pixirica-açu”. It is an exuberant and woody shrub that reaches up to 4.5 m in height and forms dense populations in the physiognomies of wet and flooded fields, riparian forest, and inner edge of the forests because it is highly tolerant to shaded environments (Romero and Martins 2002; Durigan et al. 2004; Wanderley et al. 2009). In the state of São Paulo, *M. chamissois* is an exclusive species of flooded areas and occurs along the Cerrado watercourses areas, which present distinct plant species composition and structural characteristics from the adjacent vegetation (Braga et al., 2001)

The *Miconia* species samples were identified on the basis of identification keys and the specialized literature (Romero and Martins, 2002; Souza, 2012; Wanderley et al., 2009; Zappi et al., 2015); the procedures that have already been established in the Angiosperms system were followed. The voucher specimens of the cataloged plants were confirmed and registered at the Herbarium of the Department of Biology, Laboratory of Plant Systematics, Faculty of Philosophy, Sciences, and Letters of Ribeirão Preto, University of São Paulo, Brazil (Herbarium, SPFR), under the number SPFR 16259 for *M. albicans* and SPFR 16260 for *M. chamissois*.

### Signal Collection

#### Signal Collection

The bioelectrical signals were collected by using healthy and mature leaves from the third node of the branches of adult plants, where non-invasive gold-plated polarized surface electrodes (Neurosoft 0.85 Ihm) were fixed. The electrodes were fixed with conductive gel (Carbogel EEG gel) and adhesive tape. This system kept the pressure constant and prevented the electrodes from moving on the surface of the leaves while the signal was being acquired. Two electrodes were used on both the upper and lower sides of the leaves, and a ground reference electrode was fixed in the soil, close to the sampled plant (Figure 1).

**Fig. 1.**
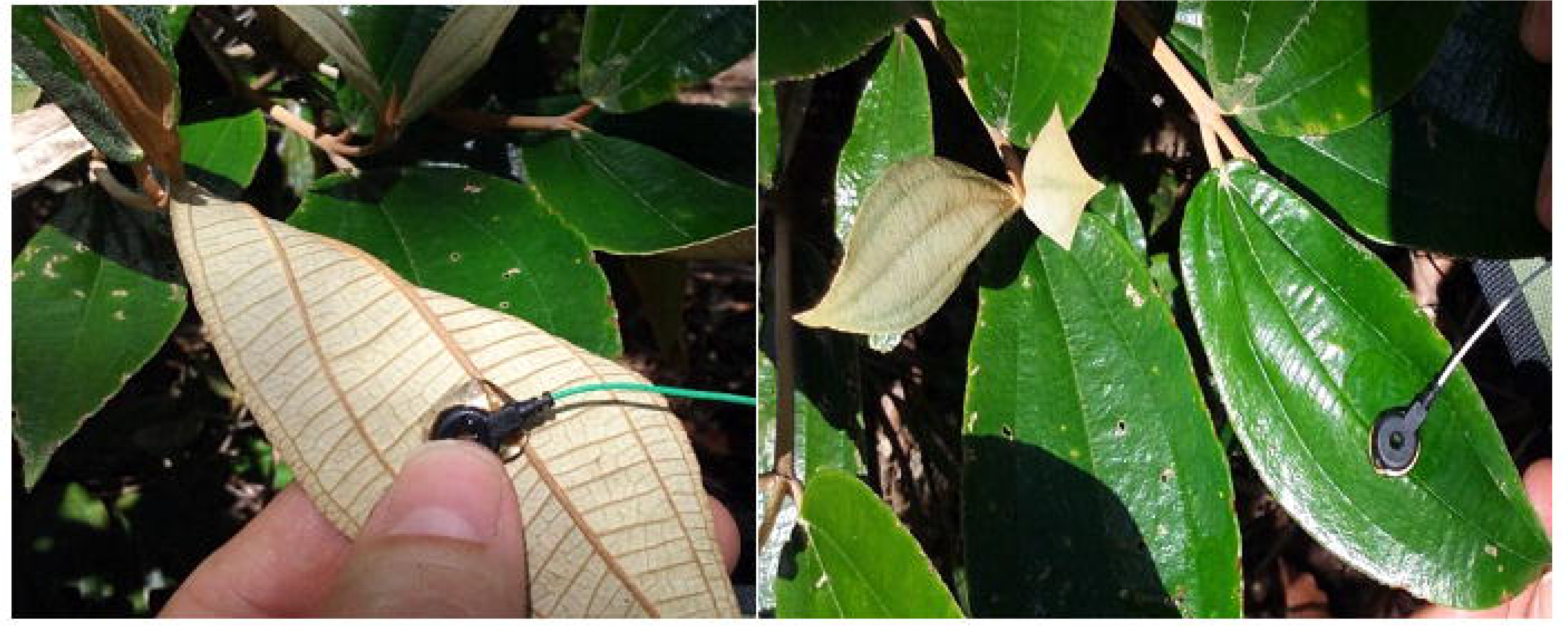
Gold electrode placement with Carbogel EEG gel on plant leaves.

The signal was sampled at 200 Hz, with two filters (High Pass and Notch), to eliminate external noise and noise from the equipment itself. Then, the signal was amplified by a factor of 10^4^ and sent wirelessly via the Bluetooth protocol to the software developed by the Laboratory of Computation and Applied Physics (LAFAC), according to Cabral et al. (2011).

The bioelectrical signal of the samples was obtained early in the morning to avoid physiological variability and depression of liquid photosynthesis around noon, when radiation becomes stronger, heat intensifies, air evaporative capacity increases, stomata ends close, intercellular CO_2_ concentration increases (there is a limitation of the photosynthetic process), and photochemical efficiency and water potential decrease (Larcher, 2006). To reduce the effects of the individual, short-period measurements (1 min) were taken and consisted in recording of 12,000 electrical signal amplitudes. Several measurements were conducted and stored; the measurements with less environmental interference (winds and sounds) were chosen as representative of the individual. The acquired signals were in the time domain, expressed in time (s) and amplitude (mV), and in the frequency domain, expressed in magnitude (Db) and frequency (Hz), and they constituted a numerical database that was used to characterize them. The frequency domain behavior of the bioelectrical signals was observed through the average spectra processed from the signals obtained from the time domain specimens.

### Signal Processing and pattern analysis

The PSD values of the bioelectrical signals sampled from the plants was calculated by using a computational algorithm implemented in the MatLab® environment. Several methods can be used to estimate PSD, the simplest one being the Welch approach with the periodogram method, which is used to determine the power density of the frequency components in a signal based on the Fourier-transform (FT). The Welch approach consists in estimating autocorrelation in the FT (obtained by averaging the autocorrelation of the segments of a bioelectrical signal sequence with 50% overlap), computing a power spectrum by using a fast Fourier-Transform (FFT) on each segment, and averaging these spectra (Welch, 1967; Barbe, Pintelon, and Schoukens, 2010) by considering the 95% confidence interval (Proakis, J.G.; Manolakis, 1996).

The PSD values obtained from the plant’s bioelectrical signal were plotted, and the raw data were divided into classes representing each plant specimen. The PSD raw data were labeled as Mad for PSD obtained for *M. albicans* growing in dry area, Maw for *M. albicans* growing in the wet area, and Mc for *M. chamissois*. Next, 50% of the PSD raw data of each class was used to train a random forest algorithm implemented in python by using the h2o library with 50 trees. The remaining 50% of the PSD data was used to test if the PSD raw data from each specimen were statistically separable. The statistical separability of the PDS raw data was analyzed by using Mean Squared Error (MSE), which represents the difference between the original and the predicted values extracted by squaring the average difference over the dataset, to obtain the Root Mean Squared Error RMSE, which is the error rate by the square root of MSE, and the Confusion Matrix (CM). All these statistical metrics were obtained from the algorithm that was developed by using the h2o python library.

The h2o python library is an open-source machine-learning tool that implements advanced algorithms such as deep learning, boosting, and bagging ensembles, among others. The h2o library can handle big data rows in-memory by using in-memory compression, even in the case of a small cluster. The algorithm that was implemented by using the h2o python library was employed to verify whether the PSD values obtained for the bioelectrical signals of plant specimens belonging to different species and collected from different areas were statically separable. The PSD features calculated from the bioelectrical signals of plants were used to prepare an input feature vector to train and to validate machine learning with random forest algorithm as illustrated in Figure 2.

**Fig. 2.**
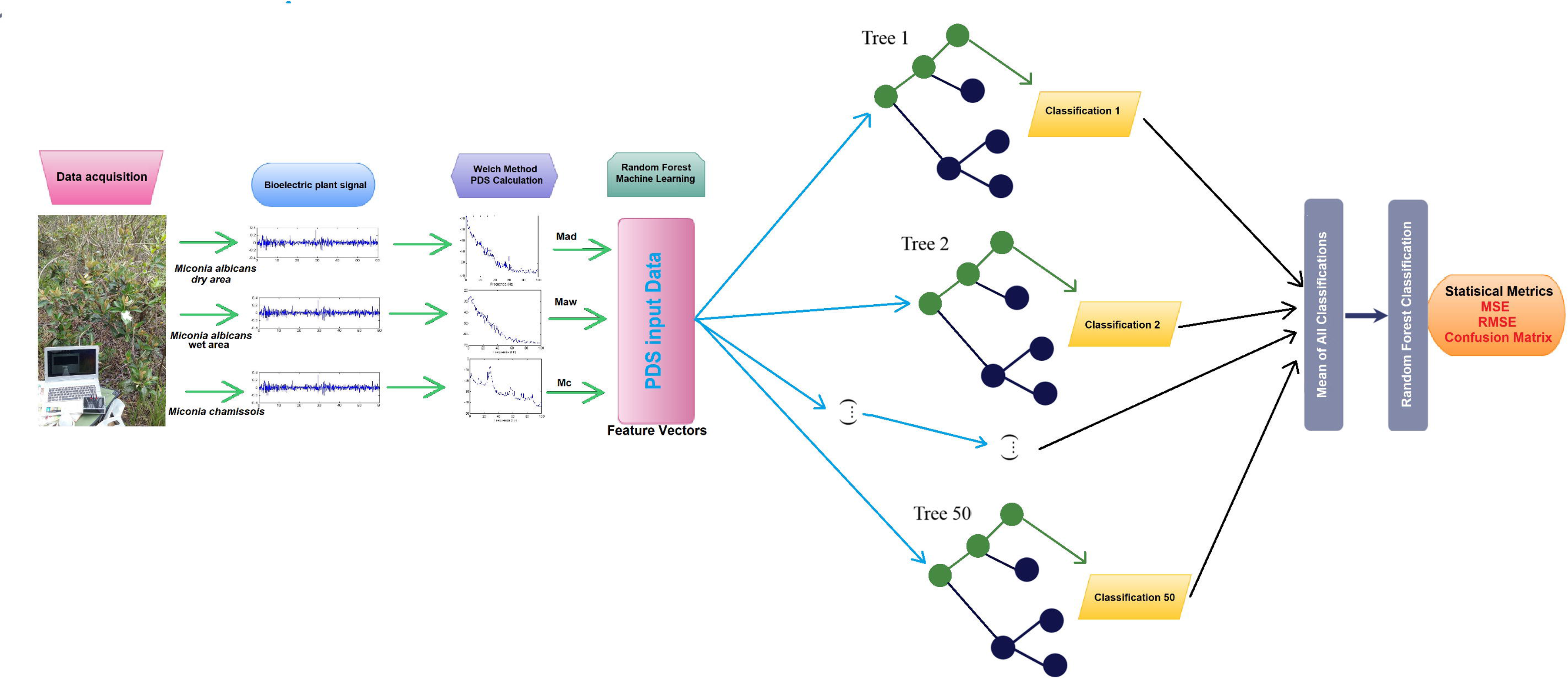
Methodology for pattern analysis with the random forest algorithm.

The feature vector was prepared by employing seven components or variables obtained from the PSD values of seven individuals of a given species. Each feature vector component was assembled with spectral points; i.e., magnitude (dB) versus frequency (Hz) point obtained from PDS as illustrated in Figure 3. PSD has 1024 spectral points, so each specimen has 1024 points to represent its spectral species pattern. In this way, 1024 vector features were generated for each specimen of a given species. On the basis of this approach, the classification algorithm was trained and tested by using 18,450 feature vectors, which amounted to 129,150 variables. Fifty percent of the total dataset was used to train the machine learning, and 50% of the data set was used in the cross-validation test based on the Confusion Matrix (CM) method (Figure 3).

**Fig. 3.**
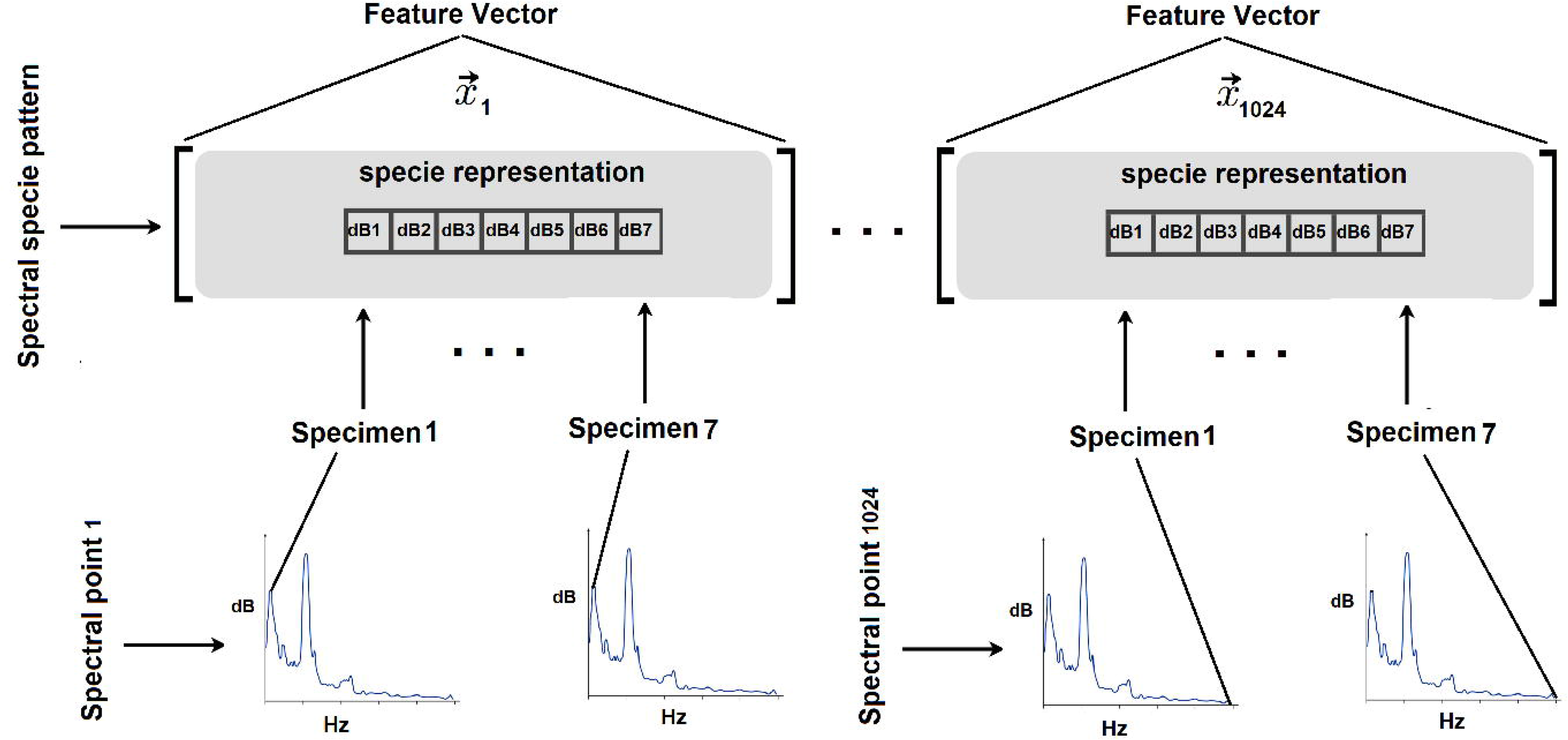
Feature vector generated with spectral points representing each species.

## Results and discussion

### Study Areas

The Brazilian Cerrado is the second largest biome in Brazil: it covers 20% or 204 million hectares of the Brazilian territory; it is situated between the Amazon and Atlantic Forests (without physical barriers); and it forms a large corridor of modest-looking xeromorphic vegetation (Oliveira and Marquis, 2002). The smaller size of the Cerrado vegetation can be explained by the fact that it resembles an inverted forest, where only about one third of the plant structure is on the soil surface. It is home to many Brazilian water springs; that is, it shelters the springs of important watersheds taking water to the Amazon Forest, the Atlantic Forest, Pantanal, and Caatinga.

Cerrado is characterized by well-defined climatic seasonality in the dry and rainy seasons, and it is considered one of the 36 hotspots of preservation. It concentrates more than 15% of all the biodiversity known in the world, and endemic species are abundant and coexist in 25 environments (Ramírez, Pringle, and Wantzen, 2008). However, 49% of the region occupied by the biome has been deforested and converted to pastures, crops, hydroelectric dams, mines, and urban areas, and only 8% of the native vegetation is protected. Apart from being a very diverse and heterogeneous system, Cerrado is very dependent on and sensitive to what happens in its surroundings. The great biodiversity of the Cerrado biome is believed to be linked to the diversity of the existing environments. Studies have tried to relate the physiognomies of the plants to environmental factors and have generally focused on associations with climate, soil classes, and fertility. Nevertheless, new insights into how living beings perceive and respond to stimuli from the environment are necessary (Gottsberger, 2006).

Biotic and abiotic interactions and the functional diversity of species over time generate an environmental gradient that acts as a filter that can shape the structure of the community in the Cerrado biome, resembling what happens in other similar biomes on the planet (Stubbs and Baston, 2004; Michalet et al., 2015; Soliveres, Smit, and Maestre, 2015). Microclimates are also relevant factors in ecosystems: they can mitigate the most severe conditions and facilitate unique community interactions (Mason et al., 2010; Molina-Venegas et al., 2018).

Studies have described some of the aspects that hinder the maintenance of Cerrado native plant species outside this biome. Examples of such aspects include great root development, greater difficulty in rooting some species, and morphophysiological adaptations to the typical Cerrado climatic stress (Cornwell, Schwilk, and Ackerly, 2006; Bulleri et al., 2016). Observations of Cerrado adult plants species have also reinforced the presence of exclusive characteristics of their development, such as the presence of xylopodium, an aspect that also prevents the maintenance of Cerrado species in a greenhouse (Melo et al., 2008). Xylopodium, whose origin (stem or root) has not yet been established, demonstrates a unique strategy of underground adaptation to explore water in deep soils (Apezzato Da Gloria, 2015).

Seedlings of the species *M. albicans* and *M. chamissois* are commercially available, but care must be adapted in terms of seed propagation, slow *M. albicans* growth rate (three or four years) and moderate *M. chamissois* growth rate, root development, and strict conditions of humidity and light for each species (Kuhlmann, 2018). Bearing in mind the various aspects attributed to the development of these plants and the complex synergy of these characteristics with numerous biotic and abiotic interactions in the natural environment, we have collected bioelectrical signals *in loco*.

### Spectral analysis

Before collecting the bioelectrical signals, we tested the data acquisition system by sampling signals with open electrodes and then with electrodes placed on plant leaves. We then compared the PSD values of the signals obtained with open electrodes to the PSD values of the plant signals. Figure 4 shows that the PSD values of the signals acquired with open electrodes displayed the characteristics of a random signal with distribution resembling the distribution of a blank noise, which differed from the signal sampled with the electrode fixed in the plant. These results showed that the signal acquired from the plants were not randomly distributed in the frequency domain and were in agreement with the work of Cabral et al. 2011.

**Fig. 4.**
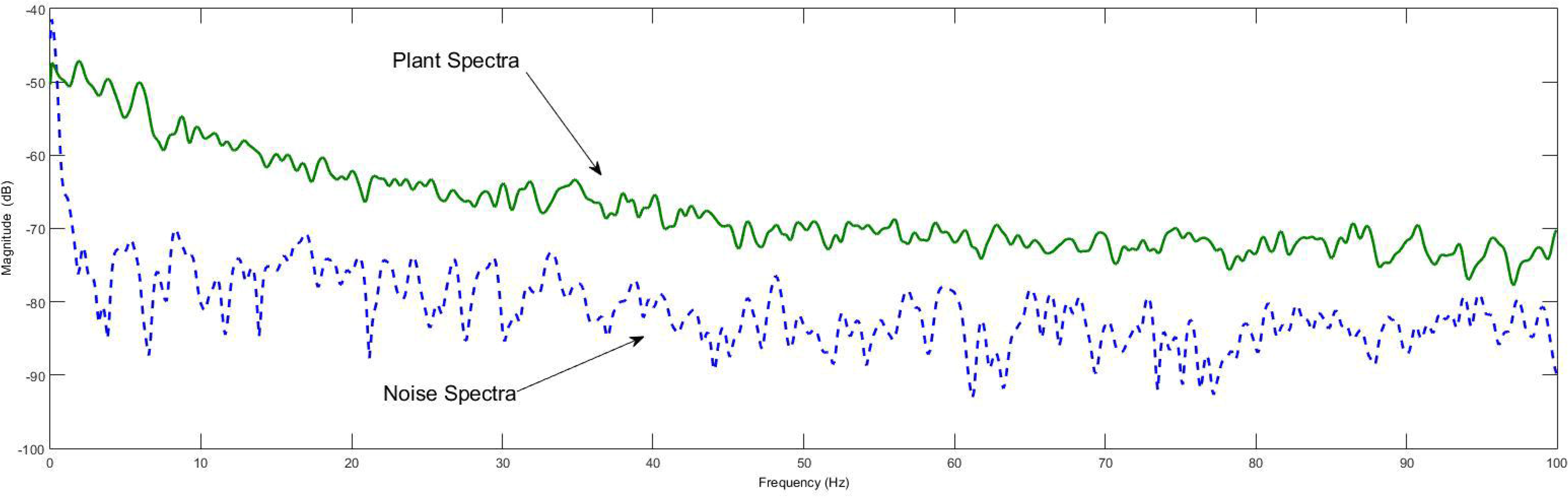
Spectral comparison between signals acquired from the plant and noises.

We carried out this procedure in all the experimental areas to certify that the acquired signals were free of interferences other than the background noise fluctuation of the open electrodes.

The bioelectrical signals obtained from the Ma (*M. albicans*) and Mc (*M. chamissois*) species in the three experimental areas revealed spectra with small peaks distributed along with the frequencies. This PSD behavior could be related to how the species interacts with environmental variables like the incidence of light, soil, and others. The plant can recognize environmental factors; for example, temperature, humidity, atmospheric pressure, and light intensity, as observed in the study of Souza et al. (2017), who even used the term “electroma” to refer to the electrical dimension of the plant as a response to the environmental variables.

The shape of the spectra obtained in the three study areas suggested the presence of a frequency pattern in the bioelectrical behavior, which was different for each of the botanical species (Figure 5) when the signals were sampled in the same season.

**Fig. 5.**
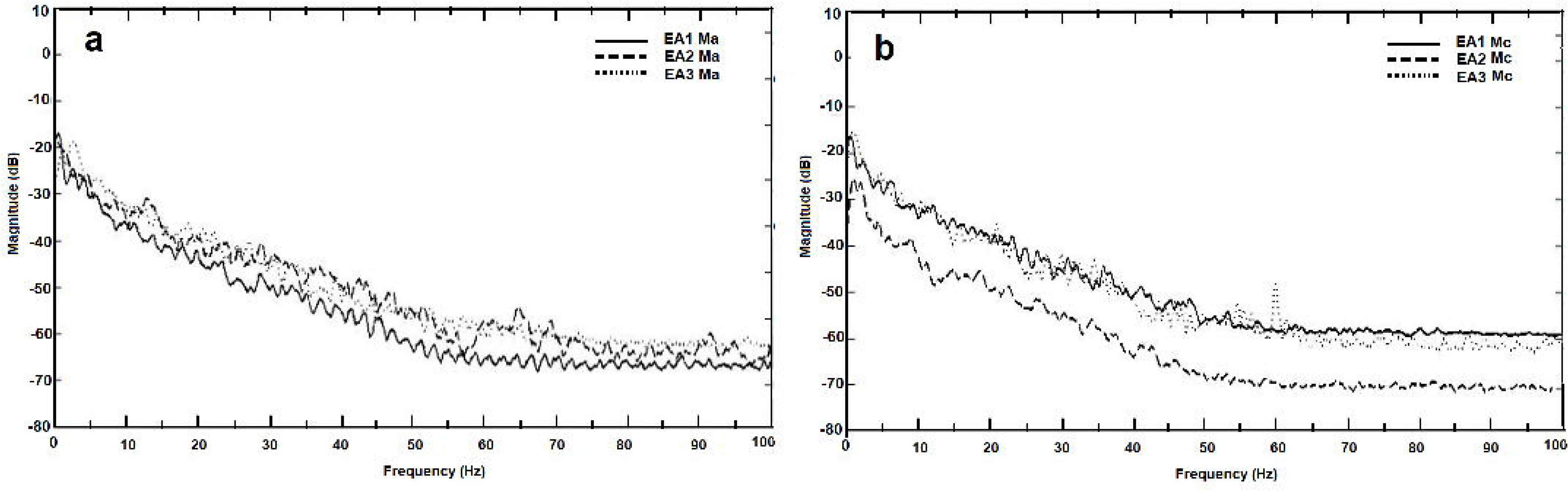
Difference in the spectral pattern of the species Ma (a) and Mc (b) for areas EA1, EA2, and EA3.

Figure 6 shows the average of the spectra recorded for the Ma and Mc bioelectrical signals collected in the three areas at different times and then averaged and plotted with their standard deviation. Given that the standard deviations did not overlap in the spectra in Figure 6, we can suggest that each species has distinct frequency behavior in the mean.

**Fig. 6.**
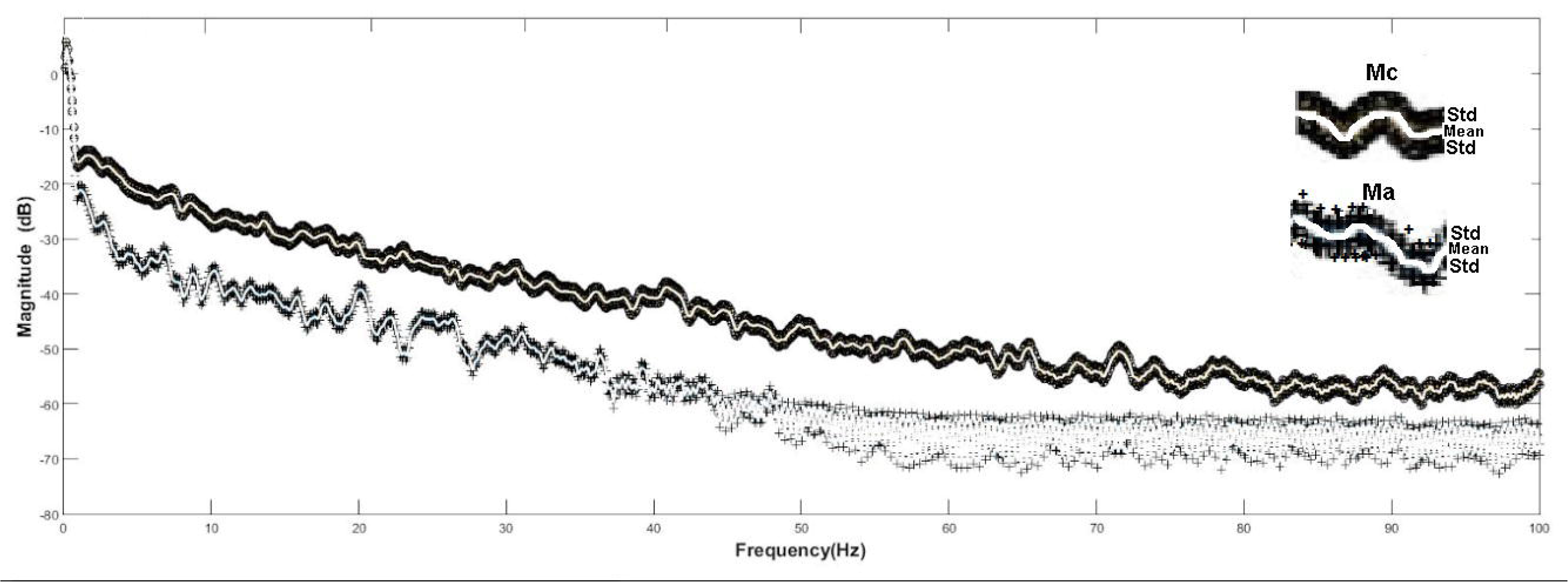
Mean spectral pattern comparison ± sd of 42 samples of Ma and Mc in different areas.

### Pattern analysis

The average spectral data were not enough to ensure that specimens had a spectral pattern, so we went back to using all the spectra of all the specimens to train an RF algorithm and then check whether there was a frequency pattern in each specimen by measuring the RF classification rates after Training. We trained the RF algorithm by using the parameters listed in Table 2.

**Table 1.** Location and climate classification according to Köpen-Geiger including coordinates and altitude (m) for year 2016 of the study areas when they were used as samples (CEPAGRI 2017).

| Study area | Climate | Coordinates | Altitude (m) | T min (°C) | T max (°C) | R min (mm) | R max (mm) | R average (mm) |
| --- | --- | --- | --- | --- | --- | --- | --- | --- |
| EA1 | Cwa | S 21°36'<br>NO 47°15' | 620 | 8 | 29.3 | 23 | 237 | 1395 |
| EA2 | Aw | S 21°33'<br>NO 47°51' | 515-650 | 10.8 | 29.6 | 26.6 | 273.6 | 1516 |
| EA3 | Cwb | S 20°08'<br>NO 47°16' | 1040 | 9.4 | 28.5 | 18 | 286.9 | 1545 |

\* minimum temperature (T min (°C)), maximum temperature (T max(°C)), minimum rainfall (R min(mm)), maximum rainfall (R max(mm)), and annual rainfall (R annual(mm)), EA1 - Campus/USP in Pirassununga, EA2 - Jataí Ecological Station (EEJ), and EA3 - Furnas do Bom Jesus State Park (PEFBJ).

**Table 2.** Python’s h2o Random Forest library model summary used to classify the PDS from the bioelectrical signals of the plants.

| Number of trees | Number of internal trees | Model size in bytes | MSE | Mean Per-Class Error |
| --- | --- | --- | --- | --- |
| 50 | 50 | 111239 | 0.019 | 0.000406 |

Table 3 shows the CM for classification of the feature vector representing the spectral pattern of Ma and Mc by the RF classifier. The CM refers to the output of the algorithm when 9217 individual PSD samples of the bioelectrical signals from the plants were presented. The data indicated that only 4 PSD values were classified incorrectly, which led to an error rate of 0.0004, hence a very small error rate. Additionally, the spectral average was not used in the RF experiments.

**Table 3.**
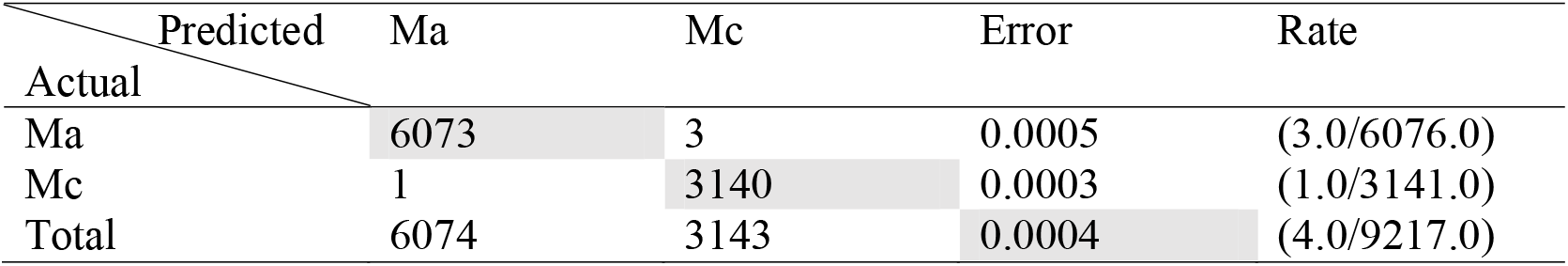
Confusion Matrix for cross validation data.

Applying machine learning in plant classification effectively helps to discover existing patterns in the various characteristics of plants. In this sense, this study agreed with other literature results based on the use of machine learning as a tool to recognize patterns in plants (see the work of Gokhale et al., 2020). Regarding the existence of plant bioelectrical patterns that can be classified with machine learning, the results of this article agreed with the results obtained by Nambo et al. (2018), who used machine learning to show the existence of bioelectrical patterns in living plants that are sensitive to environmental variables.

Finally, analysis of the mean of the spectra considering their standard deviation and of the results obtained with machine learning indicated bioelectrical characteristics that were specific to the species. This is important because bioelectricity could become a new tool to monitor and to characterize plants *in loco*.

### Conclusion

We have confirmed the hypothesis that plant bioelectrical signals can be collected *in loco*. The methodology that we used to acquire the bioelectrical signals revealed that the plant electrical signals differ from the noise pattern that could interface with the signal, and the frequency spectrum showed a characteristic average pattern for each species.

Additionally, the developed experiments generated an important database of samples of bioelectrical amplitudes for analysis in the time and frequency domain.

The frequency domain analyses based on the PSD and machine learning techniques suggested the existence of a bioelectrical pattern that is characteristic of each species, *M. albicans* and *M. chamissois*, even though the data were acquired in different Cerrado physiognomies. Comparison of the oscillatory behavior of the species left no doubt about their bioelectrical characteristics, the feasibility of collecting the bioelectrical signals, the presence of a bioelectrical pattern in *Miconia*, and the influence of environmental factors on them.

From the point of view of experimental botany, new concepts and new questions must be formulated to advance understanding of the communicative nature of plants and their interaction with the environment (Baluška and Mancuso, 2009). The results of this on-site technique represent a new methodology to acquire noninvasive information, that can be associated with other physiological, chemical, and ecological aspects of plants.

## Acknowledgements

The authors acknowledge the grants provided by Conselho Nacional de Desenvolvimento Científico e Tecnológico (CNPq) and Fundação de Amparo à Pesquisa do Estado de São Paulo (FAPESP) (Universal Program 441335/2014-4 and #2016/10313-4, respectively), and the financial support of Coordenação de Aperfeiçoamento de Pessoal de Nível Superior, Brazil (CAPES, Finance Code 001). VMM Gimenez thanks CAPES for a fellowship, and PM Pauletti and EJX Costa thank CNPq for fellowships.

## Author’s Contributions

Valéria M M Gimenez: Methodology, Investigation, Writing;

Patricia M Paulleti: Visualization, Investigation, Writing-Review & Editing, Funding acquisition;

Ana C S Silva: Formal analysis, Review & Editing;

Ernane J X Costa: Conceptualization, Supervision, Writing-Original draft preparation, Methodology, Funding acquisition;

## Data Availability

The raw data of the PDS spectra used during the study are available from the corresponding author by request.

## Abbreviations

AP: action potential
CM: Confusion Matrix
EA1: experimental area 1, in Pirassununga
EA2: experimental area 2, in Luíz Antônio
EA3: experimental area 3, in Pedregulho
LEP: local electrical potential
Ma: *Miconia albicans*
Mad: PSD of *M. albicans* of the dry area
Mc: *Miconia chamissois*
Maw: PSD of *M. albicans* of the wet area
Mc: PSD of *M. chamissois*
MSE: Mean Squared Error
SP: systemic potential
OS: oscillatory signal
PSD: power spectral density
RF: Random Forests
RMSE: Root Mean Squared Error
VP: variation potential

